# Continent-wide structure of bacterial microbiomes of European *Drosophila melanogaster* suggests host-control

**DOI:** 10.1101/527531

**Authors:** Yun Wang, Martin Kapun, Lena Waidele, Sven Kuenzel, Alan Bergland, Fabian Staubach

**Author notes:** **Competing interests** The authors declare no conflict of interest.

## Abstract

The relative importance of host-control, environmental effects, and stochasticity in the assemblage of host-associated microbiomes has been much debated. With recent sampling efforts, the underpinnings of *D*. *melanogaster’s* microbiome structure have become tractable on larger spatial scales. We analyzed the microbiome among fly populations that were sampled across Europe by the *European Drosophila Population Genomics Consortium* (DrosEU). We combined environmental data on climate and food-substrate, dense genomic data on host population structure, and microbiome profiling. Food-substrate, temperature, and host population-structure correlated with microbiome-structure. The microbes, whose abundance was co-structured with host populations, also differed in abundance between flies and their substrate in an independent survey, suggesting host-control. Patterns of enrichment and depletion of microbes between host and substrate were consistent with a model of host-control, where the host manipulates its microbiome for its benefit. Putative host-control was bacterial strain specific, supporting recent evidence for high specificity of *D*. *melanogaster-*microbe interaction.

## Introduction

Species interactions, such as the interactions between microbes and their hosts are expected to be frequently driven by competition for resources that results in evolutionary conflict (Queller and Strassmann, 2018). In particular, horizontally transmitted microbes can leave an exploited host and move on to the next host, favoring conflict (Ebert, 2013a). Nonetheless, microbes that benefit higher organisms are prevalent (e.g. Jaenike et al., 2010; Ankrah and Douglas, 2018; Bang et al., 2018). The frequent benefits that hosts derive from their microbiome are thought to result from strong selection pressure on the host to evolve selectivity towards beneficial microbes (Foster and Wenseleers, 2006; Schluter and Foster, 2012). This selectivity is also termed host-control. However, evidence for strong host-control over microbiome is limited for many organisms and its importance compared to environmental factors and stochastic processes in shaping host-associated microbiomes is debated.

Recent data across a variety of hosts, including mammals and insects, suggest that host-control might not be a major driver of microbiome structure. First, environmental factors like host-diet often have a dominant effect on microbiome composition (Chandler et al., 2011; Wu et al., 2011; Staubach et al., 2013; Wang et al., 2014; Waidele et al., 2017; Rothschild et al., 2018). Second, the discovery of substantial variation of the microbiome between individuals of the same species (Ley et al., 2008; The Human Microbiome Project Consortium et al., 2012; Linnenbrink et al., 2013; Wang and Staubach, 2018) is also difficult to explain under strong host-control. To explain this variation, stochastic, ecologically neutral processes have moved into focus. These processes comprise ecological drift, dispersal, and colonization history (Hubbell, 2001). A dominance of neutral microbiome structuring principles appears plausible for diverse species (Sieber et al., 2018), including *Drosophila* (Adair et al., 2018).

Without strong host control it is difficult to understand why higher organisms receive benefits from their microbiome so frequently. This is because host-control is supposed to be an important driver of the evolution of microbiome derived benefits in a setting that would otherwise frequently result in conflict between host and microbe (Foster and Wenseleers, 2006; Schluter and Foster, 2012; Ebert, 2013b; Queller and Strassmann, 2018). A model that resolves this contradiction has recently been proposed by Foster et al. (2017). In their ‘ecosystem on a leash’ model, the microbiome behaves similar to an ecosystem that is mainly shaped by microbe-microbe interactions. Host-control (the ‘leash’) acts only on a small subset of microbes that affect host fitness. Targeted fostering and exclusion of a relatively small set of microbes in the microbiome might suffice to favor beneficial microbiome function (Schluter and Foster, 2012; Agler et al., 2016; Foster et al., 2017).

Low but effective levels of host-control as proposed by the ‘ecosystem on a leash’-model, also holds promise to help us to better understand the interaction of *Drosophila* with its microbiome. In *D*. *melanogaster*, the microbiome has well documented effects on somatic growth and reproductive output (Storelli et al., 2011; Shin et al., 2011; Téfit and Leulier, 2017; Sannino et al., 2018; Sommer and Newell, 2018). The effects of the microbiome on these traits is caused, at least in part, by an increase in the efficiency of nutrient acquisition by the host in the presence of specific microbial taxa. For instance, reproductive (Pais et al., 2018) and nutritional (Sannino et al., 2018) benefits as well as protection from pathogens (Shin et al., 2011) can be derived from members of the family Acetobacteraceae. This family dominates the natural microbiome (Corby-Harris et al., 2007; Cox and Gilmore, 2007; Chandler et al., 2011; Barata et al., 2012; Staubach et al., 2013; Adair et al., 2018; Walters et al., 2018). However, the role of host-control in the prevalence of these potentially beneficial bacteria is unclear because the ability of *D*. *melanogaster* to shape its associated microbiome might be limited (Wong et al., 2013; Blum et al., 2013; Broderick et al., 2014). Instead, probabilistic processes contribute to gut colonization (Obadia et al., 2017) and community structure in natural populations can be explained to a large extent by neutral ecological mechanisms (Adair et al., 2018). As in mammals and other organisms, environmental factors, such as the time of collection (Behrman et al., 2018; Adair et al., 2018) or diet (Chandler et al., 2011; Staubach et al., 2013; Erkosar et al., 2018; Wang and Staubach, 2018) have a strong effect on the *Drosophila* microbiome. The food-substrate that flies live on can be a more important driver of adaptation for the microbes than the host environment in *D*. *melanogaster* (Martino et al., 2018).

On the other hand, *D*. *melanogaster* microbiome structure is associated with host genotype (Unckless et al., 2015; Chaston et al., 2016; Behrman et al., 2018), indicating genotype dependent host-control. Microbial communities within flies differ from that in their stool (Fink et al., 2013), suggesting a selection process inside the fly. Further evidence that *D*. *melanogaster* exerts control over its microbiome comes from a recent study, which found that *D*. *melanogaster* larvae potentially foster *Lactobacillus plantarum* via excretions (Storelli et al., 2018). Host-control by *D*. *melanogaster* can be highly specific and fine-tuned. Pais et al. (2018) showed that *Acetobacter thailandicus* can persist in the gut of *D*. *melanogaster*, can be dispersed by the host, and provide a fitness benefit to the host, while closely related *Acetobacter* strains cannot. In lab-reared flies, dysregulation of antimicrobial effectors leads to highly specific changes in microbiome composition; Intriguingly, these changes in community composition can preferentially select for non-pathogenic taxa over pathogenic ones, despite a close phylogenetic relationship between these microbial species (Ryu et al., 2008).

Host-control of the microbiome that increases host fitness is a key parameter of the ‘ecosystem on a leash’-model. The evidence for host-control in *D*. *melanogaster* suggests that this model might help us to understand the prevalence of benefits that *D*. *melanogaster* derives from its microbiome in the face of horizontal transmission, high intra-specific variation of microbiomes, and strong environmental effects. Given that the model was originally developed with the mammalian microbiome in mind (Foster et al., 2017), its applicability to *D*. *melanogaster* would create a common framework to understand mammalian and *D*. *melanogaster* microbiomes. As a consequence, results could become more transferable between the systems. Obviously, this is highly desirable because *D*. *melanogaster* is one of the best developed model systems in biology with many advantages over mammalian models. Together with the relatively simple microbiome (Erkosar et al., 2013) that is dominated by bacteria (Kapun et al., 2018) it holds promise for unraveling the driving forces of host-microbiome interactions (Erkosar et al., 2013; Douglas, 2018).

What is currently missing to assess whether the ‘ecosystem on a leash’ model is applicable to *D*. *melanogaster* is more information on the extent of host-control on larger ecological scales and under natural conditions. These are the conditions where potential host-control has originated by evolutionary processes. If host-control is important for *D*. *melanogaster*, imprints of host-control on the microbiome should also become apparent in a natural setting. Such imprints could be reflected by co-structure of host population genetic variation and the microbiome because host-control in *D*. *melanogaster* varies between natural populations (Behrman et al., 2018; Walters et al., 2018) and depends on host genotype (Unckless et al., 2015; Chaston et al., 2016; Pais et al., 2018). Furthermore, if the natural *D*. *melanogaster* microbiome is subject to host-control, it should show general properties of host-associated microbiomes, for example elevated levels of 16S gene copies. Finally, microbes that are subject to host-control should differ in abundance between the host and its environment because the effects of host-control outside the host should be smaller or absent.

In order to test these basic predictions and further assess the role of host-control in natural *D*. *melanogaster* populations, we profiled the bacterial microbiome, in 50 samples across Europe, using 16S rRNA gene sequencing (Figure 1 and Table S1). The sampling range covered different climates and allowed us to address the effect of environmental factors on the microbiome. We combined the 16S profiles with population level allele frequency data for more than 20,000 neutral SNPs to test for co-structure of the microbiome with host genetic variation. For further exploration of potential host-control, we tested whether microbiomes showed typical properties of host-associated communities, specifically in terms of increased 16S gene copy number. Finally, we identified bacterial taxa that correlated with host population structure. These taxa were analyzed in an independent survey, comparing fly-associated microbiomes to that of their substrate, to test whether these taxa show different abundance that indicates of host-control.

**Figure 1.**
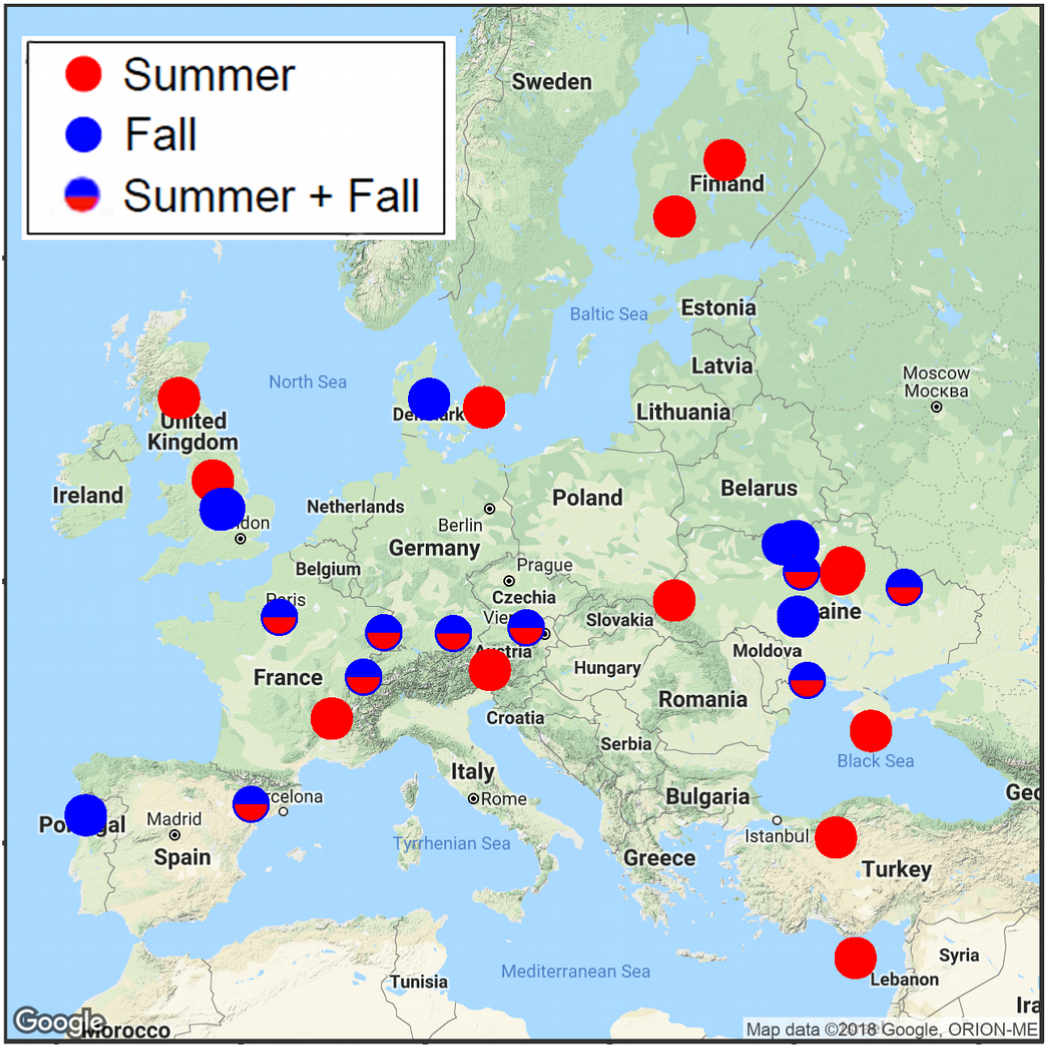
Overview of sampling locations. The map shows the geographic locations of 50 samples for bacterial community analysis in the 2014 DrosEU dataset. The color of the circles indicates the sampling season for each location.

## Results

We analyzed a total of 5,217,762 16S rRNA reads after quality filtering (Table S2). 2,672,402 *Wolbachia* sequences were removed. In order to make the diversity assessment comparable between samples, we rarefied (subsampled) the number of sequences per sample to 5,135 per sample for the continental range analysis and to 898 sequences per sample for the fly versus substrate survey. We grouped the sequences into 100% identity Operational Taxonomic Units (OTUs) for high resolution analysis, unless otherwise noted. We chose 100% identity to resolve strain level differences because the interaction with the fly host may differ for closely related bacteria (Ryu et al., 2008; Pais et al., 2018). Please note that the sequences were rigorously quality filtered and sequencing errors were removed (see Materials and Methods).

### Community composition and diversity

The *Drosophila* microbiome across Europe was dominated by acetic acid bacteria (Acetobacteraceae 63.6%, Figure S1). The three most common genera were *Acetobacter* (26.1%), *Gluconobacter* (17.4%), and *Commensalibacter* (15.4%). Enterobacteriaceae were also common (15.2%). Shannon diversity was 2.61 +/- 0.65 (SD) at the 100% identity OTU level and 1.99 +/- 0.53 (SD) at the 97% identity OTU level. Composition and diversity of the *D*. *melanogaster* microbiome were similar to those reported previously from natural *D*. *melanogaster* isolates. For a comparison of alpha diversity between studies see Staubach et al. (2013).

### The natural *D*. *melanogaster* bacterial microbiome is structured on a continental scale

As a first step to better understand the structuring principles of the *D*. *melanogaster* microbiome, we tested whether the natural *Drosophila* associated microbiome is structured on a continental scale. The absence of continental structure would be consistent with a stochastic distribution of microbes and speak against both, host-control, as well as selection of microbial taxa by other environmental factors.

Bray-Curtis-Dissimilarities (BCD) of the bacterial communities increased rapidly with geographic distance (r = 0.196, *P =* 0.0015, Mantel-Test, Figure 2), indicating geographic structure of microbial communities associated with *D*. *melanogaster* on a continent-wide scale.

**Figure 2.**
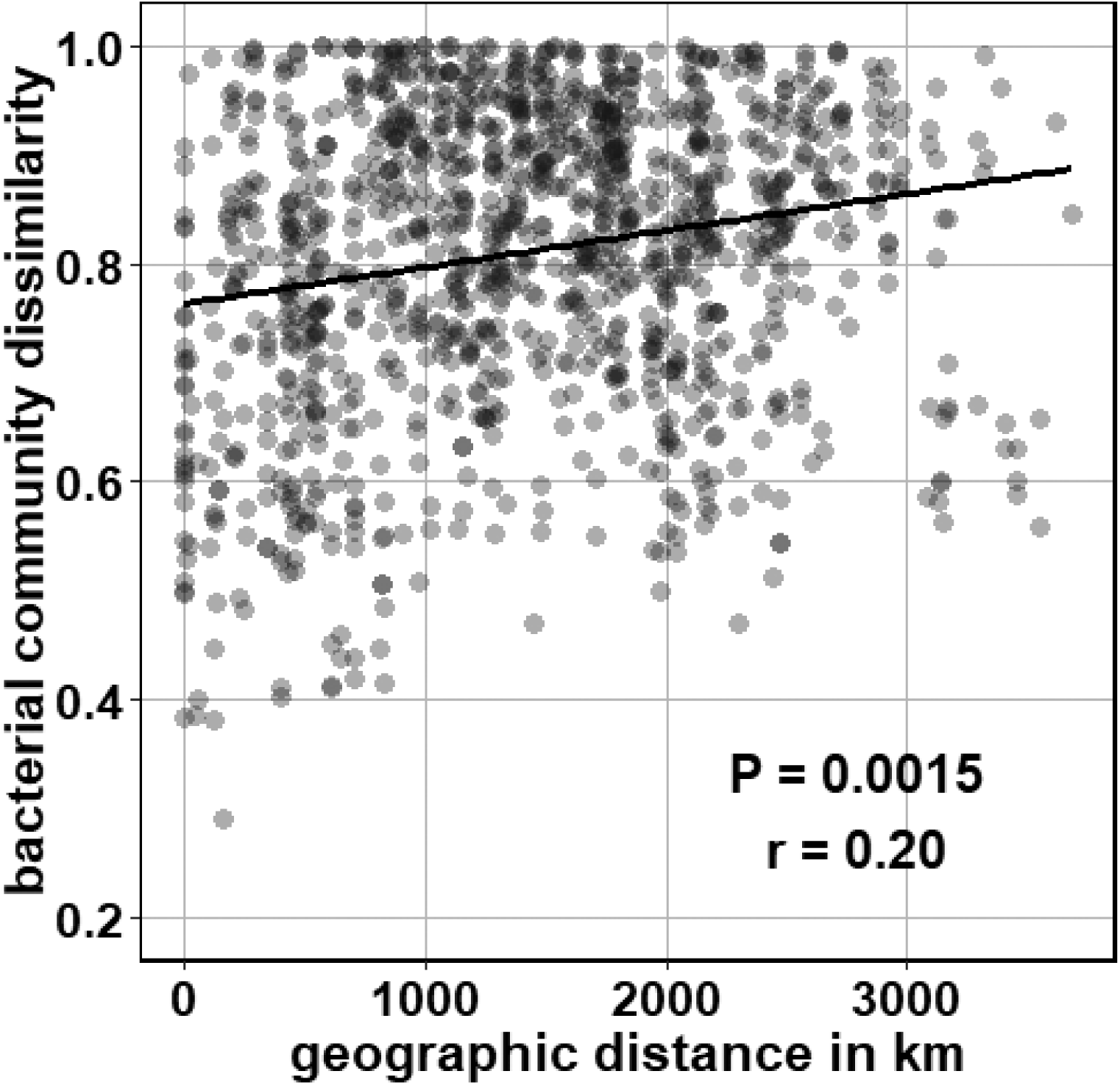
Correlation between pairwise geographic distance and BCD of the bacterial communities. P-value according to Mantel test; r is Pearson’s correlation coefficient.

### Host genetic differentiation and temperature correlate with microbiome structure

In order to identify the factors underpinning continental structure, we modeled microbiome composition in a Redundancy Analysis (RDA) framework. We selected temperature, precipitation, and substrate as candidate environmental variables that could affect microbiome structure. We chose temperature because it can affect microbial communities on geographic scales (Thompson et al., 2017). Substrate is a major determinant of natural *Drosophila* microbiomes (Chandler et al., 2011; Staubach et al., 2013; Wang and Staubach, 2018). A lack of precipitation might affect microbiome assembly by selecting for xerotolerant microbes. Because microbial communities might reflect long-term or short-term trends in temperature and precipitation, we included annual, as well as monthly means of temperature and precipitation in our model. The inclusion of monthly temperature and precipitation at the time of collection allows us to assess seasonal variation in these parameters that could affect microbiomes. Finally, we reasoned that *Drosophila* host population structure might explain interpopulation differences in microbial community composition. This would be expected if potential host-control of the microbiome varied between host populations or if microbes and fly host show patterns of co-dispersal and parallel demographic histories on a continent-wide scale.

The full model (including all factors described above) explained approximately half (46.3%) of the variance in the bacterial microbiome, indicating that the model contained factors that affect the bacterial microbiome. In order to select the most relevant explanatory variables from the full model, we employed a forward model selection approach. This resulted in host genetic differentiation (PC1 and PC2), mean annual temperature (T(y)), and substrate as relevant factors for *Drosophila* microbiome composition at the 100% OTU level (Table 1). We further reasoned that the microbiome might be structured at a higher taxonomic level. In particular, Acetobacteraceae comprise many bacteria that are dispersed by *D*. *melanogaster* and convey benefits to their hosts (Shin et al., 2011; Barata et al., 2012; Pais et al., 2018). Conversely, many Enterobacteriaceae are *Drosophila* pathogens. Susceptibility and virulence of these bacteria varies between natural host populations (Behrman et al., 2018). By applying the same model selection approach at the bacterial family level, we also identified host genetic differentiation (PC1) and annual mean temperature (T(y)) as relevant factors for microbiome composition (Table 2).

**Table 1.**
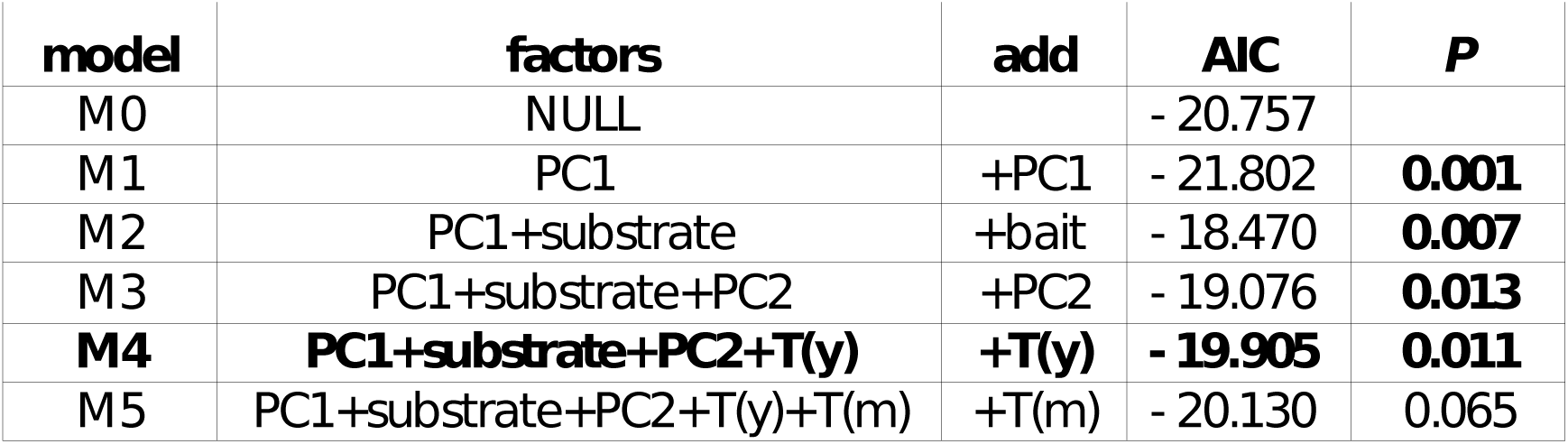
RDA model selection for factors that explain community composition at the 100% identity OTU level. The forward selection approach starts with the null model, adding the best explanatory factors one by one until adding the next factor fails to significantly improve the model. Significant p-values and the best model are in bold. P-values are based on permutation tests. PC1 = Axis 1 of host genetic variation, PC2 = Axis 1 of host genetic variation, substrate = substrate the flies were collected from, T(y) = mean annual temperature, T(m) = mean monthly temperature.

| **model** | **factors** | **add** | **AIC** | **P** |
| --- | --- | --- | --- | --- |
| M0 | NULL |  | - 20.757 |  |
| M1 | PC1 | +PC1 | - 21.802 | **0.001** |
| M2 | PC1+substrate | +bait | - 18.470 | **0.007** |
| M3 | PC1+substrate+PC2 | +PC2 | - 19.076 | **0.013** |
| **M4** | **PC1+substrate+PC2+T(y)** | **+T(y)** | **- 19.905** | **0.011** |
| M5 | PC1+substrate+PC2+T(y)+T(m) | +T(m) | - 20.130 | 0.065 |

**Table 2.** RDA model selection for factors that explain community composition at the family level. The forward selection approach starts with the null model, adding the best explanatory factors one by one until adding the next factor fails to significantly improve the explanatory power of the model. Significant p-values and the best model are in bold. P-values are based on permutation tests. PC1 = Axis 1 of host genetic variation, PC2 = Axis 1 of host genetic variation, substrate = substrate the flies were collected from, T(y) = mean annual temperature.

| **model** | **factors** | **add** | **AIC** | **P** |
| --- | --- | --- | --- | --- |
| M0 | NULL |  | - 73.782 |  |
| M1 | T(y) | +T(y) | - 76.095 | 0.002 |
| **M2** | **T(y)+PC1** | **+PC1** | **- 77.237** | **0.012** |
| M3 | T(y)+PC1+PC2 | +PC2 | - 77.488 | 0.067 |

### The abundance of OTU2 (*Commensalibacter*), and Enterobacteraceae co-vary with host population structure

We were interested to identify bacteria underlying the correlation of microbiome composition with host population structure. These bacteria might respond to potential differences in host-control between natural host populations. Therefore, we tested whether the relative abundance of OTUs correlated with host genetic variation. At the 100% identity OTU level, only the abundance of OTU2 (*Commensalibacter*, Acetobacteraceae) correlated with PC1 of host genetic variation (Figure 3, *P* = 0.00017, q = 0.0065, r = −0.52, Pearson’s Product-Moment correlation). This suggested strain level specificity of host effects on the microbiome. At the family level, Enterobacteraceae (*P* = 0.037, q = 0.17, r = 0.30, Pearson’s correlation), Leuconostocaceae (*P* = 0.047, q=0.17, r = 0.29, Pearson’s correlation), and Acetobacteraceae (*P* = 0.039, q = 0.17, r = −0.30, Pearson’s correlation) were structured according to PC1. However, when removing all sequences from OTU2, Acetobacteraceae were not significantly correlated with host population structure anymore (*P* = 0.16, q = 0.38), suggesting that OTU2 contributed significantly to the correlation. A representative sequence of OTU2 perfectly matched *Commensalibacter intestini* strain A911 (Roh et al., 2008), a previously described commensal of *D*. *melanogaster*. No individual OTU correlated with PC2 of host genetic variation.

**Figure 3.**
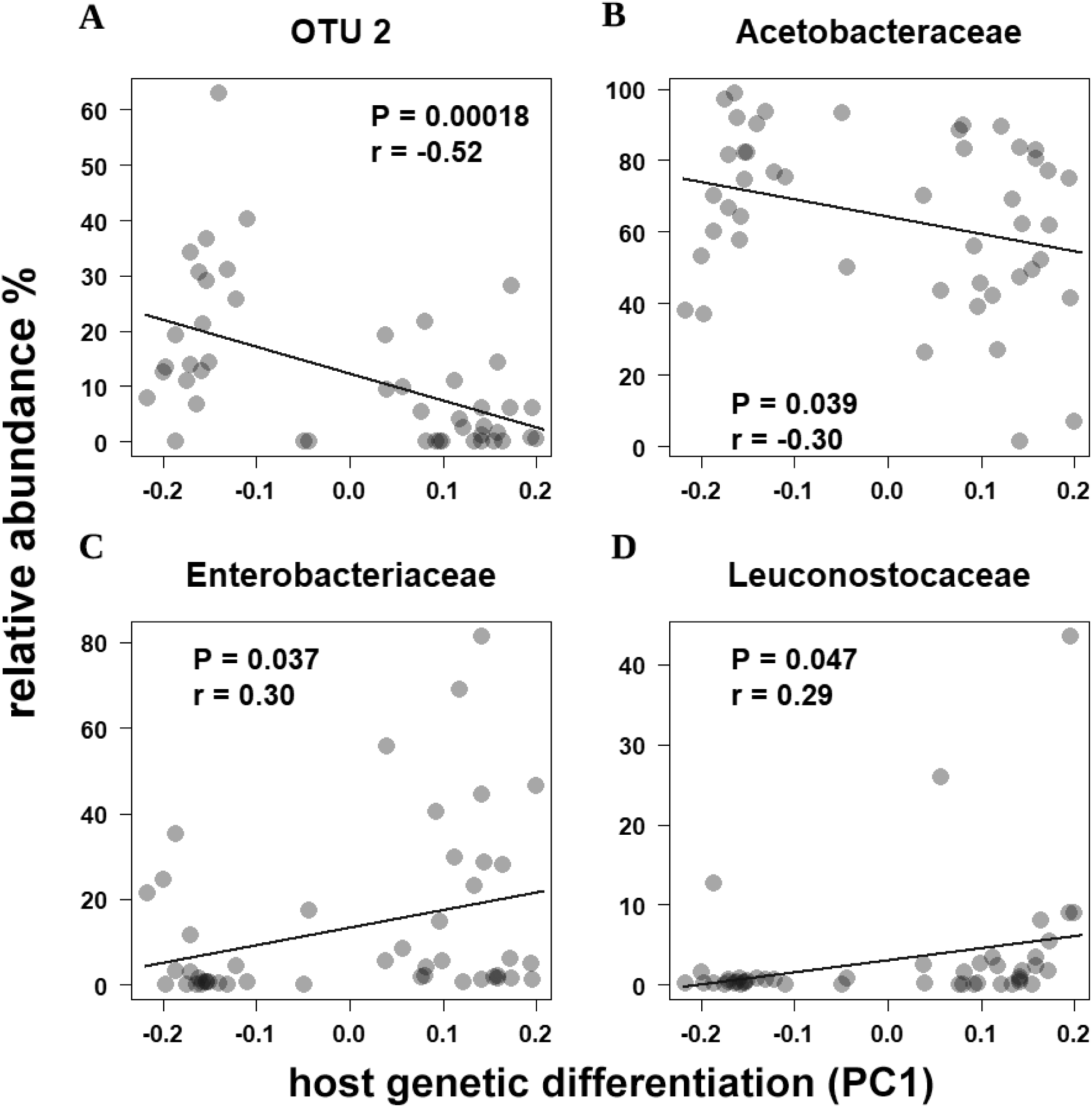
OTU2 (A) and three bacterial families (B-D) correlate with host genetic differentiation (PC1). P-value and correlation coefficient according to Pearson’s Product-Moment Correlation.

### No evidence for pronounced dispersal limitation of bacteria that correlate with host population structure

We hypothesized above that the microbiome could be affected by host-control that varies between host populations. Alternatively, a correlation of microbiomes with host population structure could result from bacteria that are dispersal limited and depend on the fly host for dispersal. If the bacteria were severely dispersal limited on a global scale, we would expect the occupied geographic range of the bacteria in question to be rather limited. However, this is not the case; the bacterial groups that are structured according to host population structure in Europe (OTU2, Enterobacteraceae, Acetobacteraceae, Leuconostocaceae) can also be found along the East Coast and on the West Coast of the USA (Figure 4). Furthermore, these bacterial groups, were also previously found in association with wild-caught *D*. *takahashii* from Hawaii, *D*. *seychellia* collected from morinda fruit on the Seychelles, cactus feeding *D*. *mojavensis* and even in mushroom feeding *Microdrosophila* (Chandler et al., 2011). A representative sequence of OTU2 matched sequences from these diverse locations and species perfectly (Chandler et al., 2011). This suggested that there is no pronounced dispersal limitation on a global scale for these bacteria and that the bacteria in question are rather cosmopolitan. Hence, a scenario, in which the bacteria are severely dispersal limited and depend on *D*. *melanogaster* dispersal on the continental scale, appears implausible.

**Figure 4.**
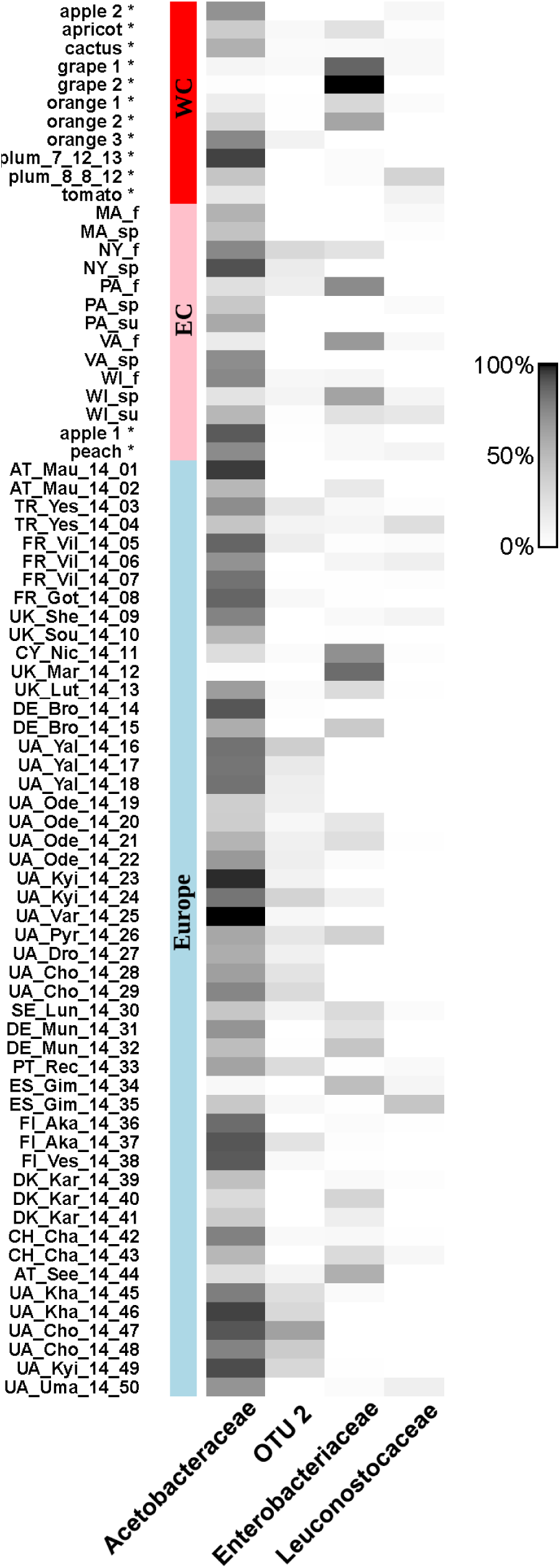
The bacteria that were structured according to host genetic variation, are common in Europe, the East Coast (EC), and the West Coast (WC) of and North America. Gray scale indicates relative abundance. Samples marked with * were described in Wang & Staubach (2018).

### 16S copy number of the natural *Drosophila* bacterial microbiome is typical for host-associated communities

Because the bacteria that are co-structured with their host populations on the continental scale are cosmopolitan, dispersal effects seemed insufficient to explain the co-structure of microbiomes and host population genetic variation. Therefore, we reasoned that host-control might contribute to the co-structure. If the *Drosophila* micrbiomes that we analyzed were subject to host-control, they should differ from environmental microbiomes. Analyzing 16S rRNA gene copy numbers can help to distinguish between environmental and host-associated microbiomes: host-associated microbiomes have increased 16S rRNA gene copy numbers (Thompson et al., 2017) when compared to environmental microbiomes. The 16S gene copy number of our samples was in the typical range of host-associated communities, and significantly higher than that of non-host associated communities (*P* < 2.2e-16, Mann–Whitney U test, Figure 5). In an independent survey, where we compared the microbiomes of flies and their immediate substrate, we also found more copies of the 16S rRNA gene in the flies than in the substrate (*P* < 0.01, Mann–Whitney U test, one-sided, Figure 5). This distinguishes the *D*. *melanogaster* microbiome from purely environmental microbiomes and supports host-related structuring.

**Figure 5.**
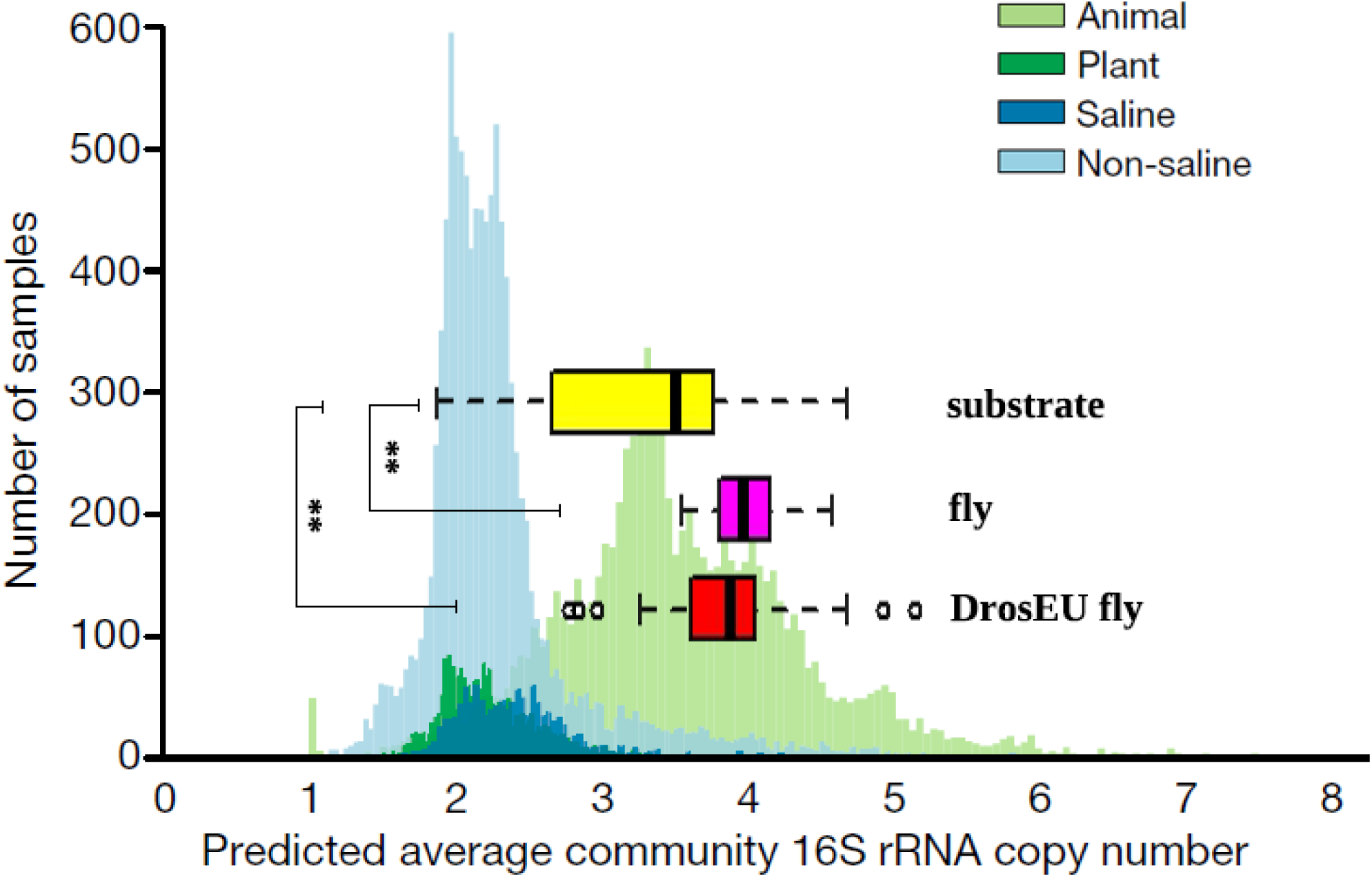
Comparison of 16S rRNA gene average copy number (ACN) between flies and substrates. ACN was higher in the fly than in the substrate communities (\*\**P* < 0.01, Mann-Whitney-U test, one-sided). Barplot (blue and green) in the background shows ACN from (Thompson et al. 2017). Fly as well as substrate samples were in the typical range of animal-associated microbiomes.

### Host-specificity of microbes that are co-structured with host population genetic variation

16S gene copy numbers suggested that the natural *D*. *melanogaster* microbiome is a typical host-associated community. This encouraged us to further explore the possibility that interactions with the host underlie the co-structure of microbiomes with host genetic variation. In order to test this, we analyzed whether the bacteria that were co-structured with host genetic variation differed in abundance between flies and their substrate. Specifically, we hypothesized that potential host-control would lead to a depletion of Enterobacteraceae in the flies because flies might avoid or reduce contact with these bacteria for their frequent pathogenicity. For example, Enterobacteraceae of the genera *Providencia, Serratia, Erwinia, Pseudomonas* are *Drosophila* pathogens. Indeed, Enterobacteraceae were more abundant in the substrate than in the flies (*P* = 0.026, paired Mann-Whitney test, one sided, Figure 6A). Furthermore, we expected to find OTU2 (*C*. *intestini*) at higher abundance in the fly than in the substrate because this OTU is a common member of the *D*. *melanogaster* associated community and contributes to healthy gut homeostasis (Ryu et al., 2008; Chandler et al., 2011). Indeed, OTU2 was enriched in flies (Figure 6B, *P* = 0.022, paired Mann-Whitney test, one-sided). Finally, we expected that Acetobacteraceae in general would be enriched in flies over substrate because this family contains several members that benefit *D*. *melanogaster* (Shin et al., 2011; Pais et al., 2018). This expectation was also confirmed (Figure 6C, *P* = 0.034, paired Mann-Whitney test, one-sided). However, when OTU2 was excluded from the analysis of Acetobacteraceae, Acetobacteraceae were not significantly enriched in flies anymore (*P* = 0.21 paired Mann-Whitney test, one sided), indicating that OTU2 contributed to family level differences. We found no difference between flies and substrate for Leuconostocaceae (*P =* 0.27, paired Mann-Whitney test, two-sided).

**Figure 6.**
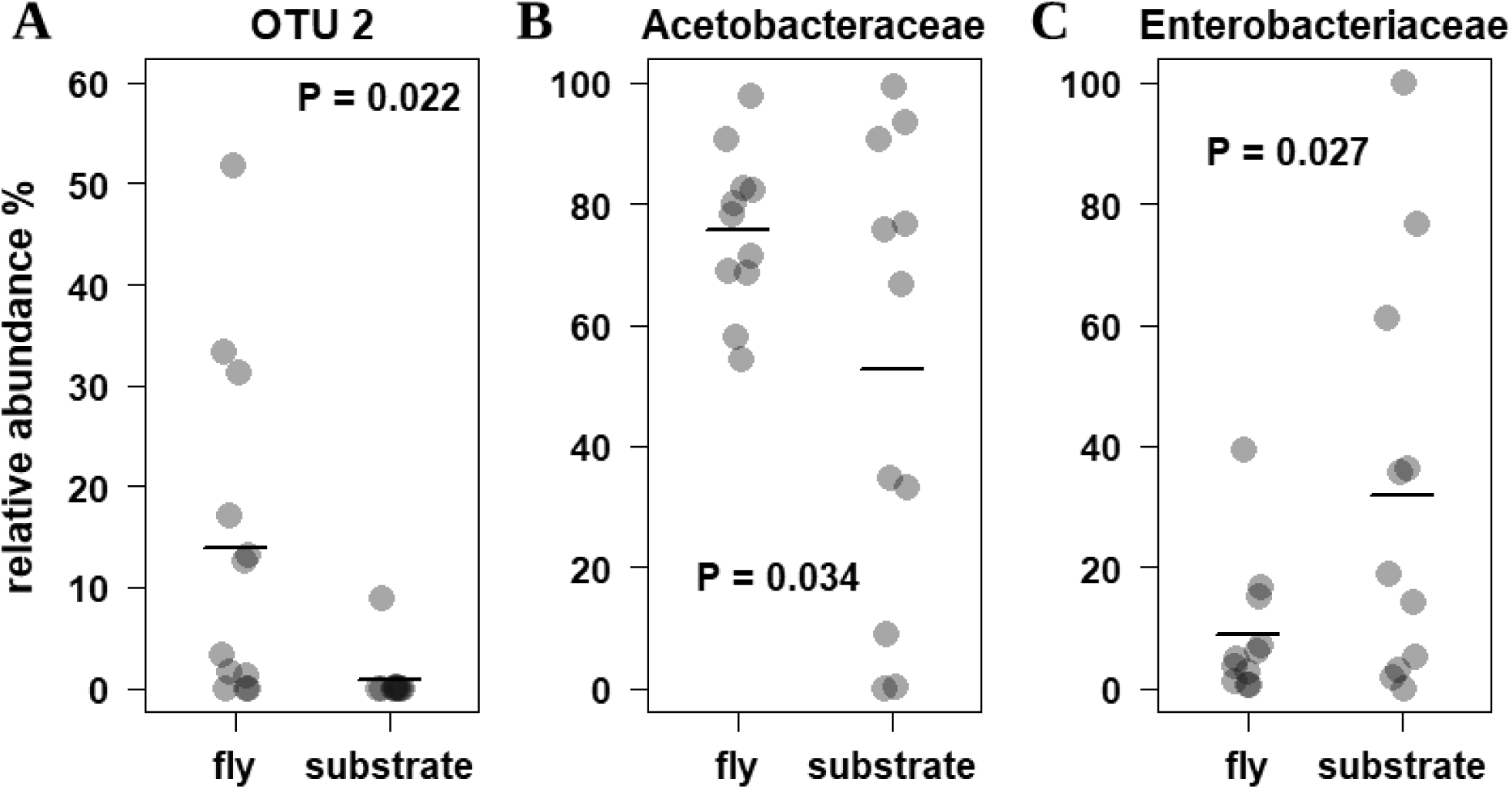
Comparison of relative abundance of (A) OTU2, (B) Acetobacteraceae and (C) Enterobacteraceae between flies and their substrate. P-values according to paired Mann-Whitney-U test, one-sided.

## Discussion

We set out to test whether there is evidence for host-control over the microbiome by *D*. *melanogaster* in a natural setting. For this purpose, we combined a comprehensive analysis of the structuring principles of the *D*. *melanogaster* microbiome on a continent-wide scale (Figure 1 and Table S1) with an independent survey comparing the microbiome of flies to that of their substrate. This resulted in several lines of evidence that support the idea that *D*. *melanogaster* exerts limited, but detectable and highly specific control over its microbiome.

### Co-structure between host genetic variation and the microbiome

The correlation of host population genetic differentiation and the differentiation of microbiomes can be interpreted as evidence for host-control. This correlation is consistent with a model, in which stronger genetic differentiation leads, on average, to larger differences in host-control, and hence host-associated microbiomes. Given ample evidence for variation in host-control between natural populations that depends on genotype (Lazzaro et al., 2008; Corby-Harris and Promislow, 2008; Behrman et al., 2018; Walters et al., 2018) this seems a reasonable model.

It appears unlikely that co-structure resulted from environmental factors that affect both, the microbiome and host genetic variation for two reasons: First, we accounted for the most plausible environmental factors that could affect microbiomes and the host at the same time in our model (food-substrate, temperature, precipitation). Second, for assessing host genetic variation, we used SNPs from small introns that are considered least affected by natural selection (Parsch et al., 2010; Lawrie et al., 2013). Therefore, it is unlikely that selection exerted by environmental factors that also affect the microbiome strongly affects these SNPs and generates co-structure.

It is similarly difficult to explain the co-structure by co-dispersal of *Drosophila* and bacteria because we found no evidence for pronounced dispersal limitation of the bacteria that co-vary with host genetic differentiation on a global scale (Figure 4). Instead, our data and previous studies suggest that these bacteria are cosmopolitan (Cox and Gilmore, 2007; Chandler et al., 2011). Taken together that environmental variation was accounted for and that we found no evidence for dispersal limitation, a role for host-control in the observed co-structure appears plausible.

### 16S copy numbers and differences between flies and substrate microbiomes support host-control

Host effects on the microbiome were further supported by 16S rRNA gene copy numbers that were in the typical range for host-associated communities and significantly different from that of non-host associated communities (Figure 5). As expected, the copy number in the substrate samples was smaller than that in fly samples. Interestingly, the copy number in substrate microbiomes was still larger than that of typical non-host associated microbiomes. This is consistent with *Drosophila* also affecting the microbiome of its immediate environment (Wong et al., 2015; Chaston et al., 2016; Storelli et al., 2018) and transforming it to appear more host-like.

Besides the increased number of 16S gene copies, host-control was evident from differences between the host microbiome and that of its substrate; three of the four bacterial groups (Acetobacteraceae, Enterobacteraceae, OTU2) that correlated with host genetic variation on a continental scale (Figure 3) also differed in abundance between flies and their substrate (Figure 6).

### Fitness effects of microbes that show evidence of host structuring support host-control

The evidence above supports host-related structuring of the microbiome that is consistent with host-control. However, the term ‘host-control’ also implies that the effects of the host on the microbiome provide some fitness benefit to the host. The bacteria that are structured in the host environment and the direction of the structuring (enrichment or depletion) suggest such fitness benefits.

It seems reasonable to assume that the reduction of Enterobacteriaceae in the fly environment (Figure 6), is likely beneficial for the flies, because this family comprises a range of the most important *D*. *melanogaster* pathogens. Examples are *Providencia* (Galac and Lazzaro, 2011), *Serratia* (Flyg et al., 1980; Lazzaro et al., 2006), *Erwinia* (Basset et al., 2000), and *Pseudomonas* (Vodovar et al., 2005). A reduction of Enterobacteriaceae in the fly gut is in line with results from Ryu et al. (2008). These authors have shown that Enterobacteriaceae, including the highly pathogenic *Erwinia carotovora carotovora-15* do not persist in the fly gut.

In contrast to Enterobacteriaceae, Acetobacteraceae were enriched in the host. This pattern was mainly driven by OTU2 (*C*. *intestini*). This OTU matches sequences from previous studies on fruit flies in the natural environment (Blast results Table S3) (Cox and Gilmore, 2007; Chandler et al., 2011; Wang and Staubach, 2018) and in the laboratory (Ryu et al., 2008). In particular, it perfectly matches *C*. *intestini strain A911* (Roh et al., 2008). This strain is sensitive to anti-microbial peptides (AMPs) (Ryu et al., 2008), and hence can be subject to host-control. In wild-type flies, it is a dominant member of the microbiome. When AMPs are misregulated it is replaced by *Gluconobacter morbifer* that has detrimental effects on flies. Thus, in the wild-type gut environment, *C*. *intestini strain A911* is favored by the host and has a protective function. Favoring of a protective microbe can be considered host-control.

### The specificity of host-control

*G*. *morbifer* as well as *C*. *intestini* are Acetobacteraceae, and hence relatively closely related. That flies can favor one over the other, points towards highly specific host-control in *D*. *melanogaster*. Our results suggest that host-control can also be highly specific under natural conditions; Only OTU2 (perfect sequence match with *C*. *intestini*) was strongly co-structured with host genetic variation and at the same time enriched in flies over substrate (Figure 3 and 6). The evidence for high specificity in the interaction with bacteria that we found parallels recent results from Pais et al. (2018). These authors found that *Acetobacter thailandicus* colonizes *D*. *melanogaster* and persists in the gut, while a closely related *Acetobacter* strain does not persist. High specificity also fits in with results from Adair et al. (2018), who showed that the assembly of the natural bacterial microbiome in the *D*. *melanogaster* gut can be largely explained by neutral processes, except for a specific set of bacteria. This specificity of the interaction of *D*. *melanogaster* with its microbiome is also fully compatible with recent advances in understanding the mechanistic principles of *D*. *melanogaster* immunity. A combination of highly specific regulation of the IMD pathway via different peptidoglycan recognition proteins (PGRPs) and specific regulation of the duox pathway (Ha et al., 2005; Lhocine et al., 2008; Bosco-Drayon et al., 2012; Lee et al., 2013; Guo et al., 2014; Iatsenko et al., 2016; Neyen et al., 2016) can lead to highly specific selection processes acting on bacterial communities in the fly gut.

While we found support for specific interaction with OTU2 (*C*. *intestini)*, more general mechanisms seemed to be at work for the interaction with Enterobacteriaceae. The reduction of Enterobacteraceae in the fly, when compared to the substrate was not linked to any specific OTUs from this family. Likewise, the co-structure with host genetic variation, was only apparent for the family as a whole. This family level host-control could arise in response to signals that are common to Enterobacteriaceae or from their potential pathogenicity in the sense of a danger or damage signal (Matzinger, 2002). Alternatively, the *Drosophila* gut might be just less favorable in terms of its physical condition (e.g. pH) or presence of antimicrobial agents (e.g. AMPs).

### Environmental factors and the *D*. *melanogaster* microbiome

In addition to host genetic structure, temperature as well as the substrate, the flies were collected from, correlated with microbiome structure. While the effects of substrate on the fly microbiome are well described (Chandler et al., 2011; Staubach et al., 2013; Wang and Staubach, 2018), a continental scale temperature effects on a host-associated microbiome has, to our knowledge, not been described before. Temperature affects environmental microbiomes on a global scale (Zhou et al., 2016; Thompson et al., 2017) and the effect in *Drosophila* might reflect the exposure of flies to different environmental microbiomes. Temperature dependence of the microbiome could also be the result of a temperature dependent dietary switch (Brankatschk et al., 2018); small scale structure of the food sources might allow flies to acquire selectively more plant or yeast material, which might lead to changes in microbiome composition.

While there was a significant effect of annual temperature on the microbiome, the correlation with monthly temperature at the collection date only showed a trend (*P* = 0.065). Because our seasonal sampling was relatively limited (nine locations), more data is required to address the question whether seasonal temperature changes affect the microbiome. Seasonal variation in *D*. *melanogaster* associated microbiomes has been described by Behrman et al. (2018) and correlates with differences in pathogen susceptibility of the host. This points towards the possibility that seasonal changes in the microbiome could add to seasonal selective regimes and contribute to seasonal genome variation in *D*. *melanogaster* (Bergland et al., 2014; Machado et al., 2018).

The effects of temperature on the microbiome seemed more general as no specific OTU nor family was significantly correlated with temperature variation.

## Conclusion

*D*. *melanogaster* lives in a microbe rich environment; rotting fruit. In this environment, it is essential for flies to foster beneficial microbes and avoid pathogens. Using continental scale data from natural populations, we presented evidence for specific host-control that favors a protective bacterium. This adds to the recent notion of high specificity of host-microbe interaction in *D*. *melanogaster* and shows that this specificity unfolds in an ecological and evolutionary context. At the same time, our study supports previous findings of strong environmental effects on the natural *D*. *melanogaster* microbiome. Strong environmental effects in combination with host-control of a relatively small subset of the partners is concordant with the ‘ecosystem on a leash’ model. Our results support the idea that this model might serve as a common framework to understand *D*. *melanogaster* and mammalian microbiomes. A common framework for mammalian and *D*. *melanogaster*, increases the transferability and generalizability between systems. Hence, we see a bright future for the *D*. *melanogaster* microbiome as a model for other organisms, including mammals, in host-microbiome research.

## Materials and Methods

### Fly and substrate samples

European fly samples were collected as described in Kapun et al. (2018). In short, 50 samples of *D*. *melanogaster* were collected from 31 locations across the European continent with a joint effort of European research groups (Figure 1 and Table S1). Each sample contained a pool of 33-40 wild-caught males. We used males only because only males can be reliably distinguished from sympatric *D*. *simulans*. The effects of pooling on *D*. *melanogaster* microbiome profiling were assessed in detail by Wang and Staubach (2018). In short, pooling provides a more comprehensive picture of the population microbiome than an individual fly. While differences in microbiome structure between individuals tend to even out in a pool, differences between populations be well differentiated. Because we were interested in variation between populations here, a pooling approach is well suited. All 50 samples were included for analyzing *Drosophila*-associated bacterial community composition, diversity and dispersal patterns. Because data on host genetic differentiation for samples FR_Vil_14_06, UA_Yal_14_17, and DK_Kar_14_40 was not available, these samples were excluded from the analysis of continental scale community structure. The visualization of fly samples on the map in Figure 1 was generated with the R package ‘ggmap’ (Kahle and Wickham, 2013).

Twelve samples from the East Coast of the USA were collected in the same fashion as the European samples and represent population pools of males. Seven of these samples were already analyzed in Behrman et al. (2018) (see Table S1 for details). The samples named NY and WI were described in Machado et al. (2018). However, the 16S data for these samples was generated here. Because we did not have detailed information on the substrate, these samples were collected from, we did not include them in our continental scale modeling. We used these samples for evaluating the global range of bacteria (Figure 4). For the same purpose, we included 13 fly samples from Wang and Staubach (2018) that were primarily collected at the West Coast of the USA (see Table S1 for details).

For the survey of the microbiome of flies and their substrate, pairs of pools of five flies and the corresponding substrate for a total of 24 samples were collected. The immediate substrate, on which the flies that we collected were sitting and feeding was collected with a sterile scalpel and transferred to a sterile microcentrifuge tube. The survey spanned 6 different substrates from 4 locations (Table S1).

### DNA extraction, PCR and sequencing

DNA from the DrosEU samples was extracted by standard phenol-chloroform extraction after homogenization with 3 minutes of bead beating on QIAGEN TissueLyser II as described in Kapun et al. (2018). DNA from population pools from the USA were extracted as described in Bergland et al. (2014). DNA extraction for pools of five flies and the corresponding substrate was performed using the Qiagen QIAamp DNA extraction kit (Qiagen, Carlsbad, CA) combined with bead beating in the same way as for fly samples from Wang and Staubach (2018).

Barcoded bacterial broad range primers, 515F (5’GTGCCAGCMGCCGCGGTAA3’) and 806R (5’GGACTACHVGGGTWTCTAAT3’) from Caporaso et al. (2011) were used to amplify the V4 region of the bacterial 16S rRNA gene. DNA was amplified with Phusion® Hot Start DNA Polymerase (Finnzymes, Espoo, Finland) under the following conditions: 30 sec at 98°C; 30 cycles of 9 sec at 98°C, 60 sec at 50°C and 90 sec at 72°C; final extension for 10 min at 72°C. In order to reduce PCR bias, amplification reactions were performed in duplicate and pooled. PCR products quantified on an agarose gel and pooled in equimolar amounts. Extraction control PCRs were negative and excluded. The resulting pool was gel extracted using the Qiaquick gel extraction kit (Qiagen, Carlsbad, CA) and sequenced on an illumina MiSeq sequencer reading 2 × 250bp.

### Data analysis

We analyzed sequencing data using MOTHUR v1.40.0 (Schloss et al., 2009). Main processing steps in MOTHUR included alignment of paired reads, quality filtering, removal of PCR errors, removal of chimeric sequences, subsampling (rarefication), and alpha-diversity calculations (Kozich et al., 2013). Sequences were taxonomically classified using the SILVA reference database ‘Release 132’ (Pruesse et al., 2007) as implemented in MOTHUR. A detailed step by step analysis script with all commands executed can be found in the supplementary File Script1 for full reproducibility. For studying geographical microbiome structure, OTUs were clustered at 100% identity. Only for the comparison of Shannon diversity to previous studies, we also included clustering at 97% sequence identity.

To identify factors that shape microbial communities, we applied Redundancy Analysis (RDA). Following Borcard, Gillet and Legendre (2018), OTU count data was Hellinger transformed to allow analysis in the linear RDA framework. In order to reduce the effects of rare species on RDA and assuming that ecologically relevant species should be frequent, we focused the analysis on OTUs with more than 1000 reads across samples. Our candidate explanatory variables were temperature, precipitation, substrate and host genetic differentiation. Data of annual and monthly mean temperature (BIO 1 and tmean) and precipitation (BIO 12 and prec) were downloaded from WorldClim (Fick and Hijmans, 2017, see supplementary File Script2). Host genetic differentiation was represented by the first two principle components of an allele frequency based Principle Components Analysis performed by Kapun et al. (2018). In short the data represents allele frequencies from more than 20,000 SNPs in short intronic sequences that evolve putatively neutral and best represent population structure. In order to select the variables that were most important for microbiome structure, we applied forward model selection of additive linear models. This was done with the ordistep function from the vegan R package (Oksanen et al., 2018). The Ordistep function provides a stepwise approach to select variables based on permutation P-values and Akaike’s Information Criterion (AIC).

In order to test for potential spatial autocorrelation we followed the protocol by (Borcard et al., 2018) using the dbmem function. This protocol employs eigenvector analysis to detect autocorrelation at different scales. We found no evidence for significant autocorrelation in our data (see supplementary File Script3) after removal of the continent-wide trend in species distributions that we analyzed here. All algorithms were part of the vegan (Oksanen et al., 2018) and adespatial R packages (Dray et al., 2018). Geographic distances were computed with the gdist function from the Imap R package (Wallace, 2012, see supplementary File Script4).

For the correlation of host genetic differentiation with the relative abundance of individual OTUs and bacterial families, we calculated q-values with the p.adjust function in R to account for multiple testing. Following the recommendation by Efron et al. (2007), only significant correlations (*P* < 0.05) with bacterial groups with q-values smaller than 0.2 were considered significant.

Average community 16S rRNA gene copy number (ACN) was predicted from 16S rRNA gene amplicon data using PICRUSt (Langille et al., 2013). The method for calculating ACN was adapted from Thompson et al. (2017). We first classified sequences using the Greengenes reference database and generated an OTU table in the Biom-format using the make.biom function in MOTHUR. The resulting biom-formated table served as input for the normalize_by_copy_number.py command implemented in PICRUSt. The output file is a normalized observation table. ACN for each sample were calculated as the raw sample sum divided by the normalized sample sum.

## Data availability

Raw sequence data is in the submission process to the ncbi short read archive (SRA).

## Acknowledgements

We thank the DrosEU consortium for the coordinated sampling and fruitful collaboration. We thank the Paul Schmidt (University of Pennsylvania), John Pool (University of Wisconsin), Lazzaro (Cornell University) labs, as well as the Drosophila Real Time Evolution Consortium (DrosRTEC) for samples. We thank Carsten Dormann (University of Freiburg) for statistical support. The authors acknowledge support by the state of Baden-Württemberg through bwHPC. This work was funded by a DFG grant to FS (STA1154/4-1, 408908608) and the Landesgraduiertenfoerterung of the state of Baden-Württemberg.

